# Fan cells in layer 2 of lateral entorhinal cortex are critical for episodic-like memory

**DOI:** 10.1101/543777

**Authors:** Brianna Vandrey, Derek L. F. Garden, Veronika Ambrozova, Christina McClure, Matthew F. Nolan, James A. Ainge

## Abstract

The lateral entorhinal cortex (LEC) is a critical structure for episodic memory, but the roles of discrete neuronal populations within LEC are unclear. Here, we establish an approach for selectively targeting fan cells in layer 2 (L2) of LEC. Whereas complete lesions of the LEC were previously found to abolish associative recognition memory, we find that after selective suppression of synaptic output from fan cells mice still recognise novel object-context configurations, but are impaired in recognition of novel object-place-context associations. Our experiments suggest a segregation of memory functions within LEC networks and indicate that specific inactivation of fan cells leads to behavioural deficits reminiscent of early stages of Alzheimer’s disease.

## Introduction

Episodic memory relies on the hippocampus and its surrounding cortical network. Within this network, the lateral entorhinal cortex (LEC) and hippocampus are required for episodic-like memory in rodents (Langston & Wood, 2010; Wilson et al. 2013a), and the LEC is critical for integrating spatial and contextual information about objects (Van Cauter et al. 2013; Wilson et al. 2013a; 2013b; Chao et al. 2016; Kuruvilla & Ainge, 2017). LEC neurons encode objects within an environment, spatial locations where objects were previously experienced, and generate representations of time during the encoding and retrieval of episodes (Deshmukh & Knierim, 2011; Morrissey & Takehara-Nishiuchi 2014; Montchal et al. 2018; Tsao et al. 2013; 2018; Wang et al. 2018). However, it remains unclear how specific populations of cells within the LEC contribute to the integration of episodic memory components.

Projections to the hippocampus from layer 2 (L2) of LEC are of particular interest given that this layer manifests early pathology in Alzheimer’s Disease (AD) and related animal models (Gomez-Isla et al. 1996; Stranahan & Mattson, 2010; Khan et al. 2014). These projections arise from a superficial sublayer (L2a) which contains fan cells that are immunoreactive for reelin and project to the dentate gyrus (Fujimara & Kosaka, 1996; Leitner et al. 2016). These neurons appear to have distinct roles in odor information processing by the LEC (Leitner et al. 2016). An impaired ability to associate different features in memory is characteristic of early AD (Fowler et al. 2002; O’Connell et al. 2004; Parra et al. 2009; 2010) and although correlations between these deficits and pathology of the superficial entorhinal cortex suggest contributions to episodic memory processes (Kobro-Flatmoen et al. 2016), the roles of specific cell types in associative memory are unclear.

Here, we ask if suppressing synaptic output from fan cells replicates the behavioural deficits found following complete lesions of the LEC. We show that Sim1:Cre mice, which were previously used to target reelin positive stellate cells in L2 of the medial entorhinal cortex (MEC; Sürmeli et al. 2015, Tennant et al. 2018), can be repurposed to give genetic access to reelin positive fan cells in L2 of LEC. By selectively blocking their synaptic output, we show that fan cells are critical for memory of object-place-context context associations, but are not required for novel object or object-context recognition.

## Results

### Sim1:Cre mice give genetic access to fan cells in L2 of LEC

Because the arrangement of neurons in L2 of the LEC is similar to L2 of the MEC (Supplemental Figure 1.1; Fujimara & Kosaka et al. 1996; Leitner et al. 2016), we investigated whether Sim1:Cre mice provide specific access to reelin positive fan cells in L2 of the LEC. We found that following injection of Cre-dependent AAV encoding green fluorescent protein (AAV-FLEX-GFP; Murray et al. 2011) into the superficial LEC of Sim1:Cre mice (n = 4), the majority of cells expressing GFP were located in LEC L2a (Figure 1A-B). Staining with antibodies against reelin and calbindin revealed that expression of GFP was specific to cells that were positive for reelin, but not calbindin (Figure 1C-D). A small number of cells were triple-labelled by reelin, calbindin, and GFP (Supplemental Figure 1.2). To establish whether the labelled population of neurons overlapped with the population of cells that project to the hippocampus, we injected a further cohort of Sim1:Cre mice (n = 3) with AAV-FLEX-GFP into superficial LEC and a retrograde tracer into the dorsal dentate gyrus. The majority of neurons expressing GFP co-localised with neurons labelled by the retrograde tracer (Figure 1E-F). Therefore, the Sim1:Cre mouse provides genetic access to a population of neurons in LEC L2 which are positive for reelin and project to the hippocampus.

**Figure 1:**
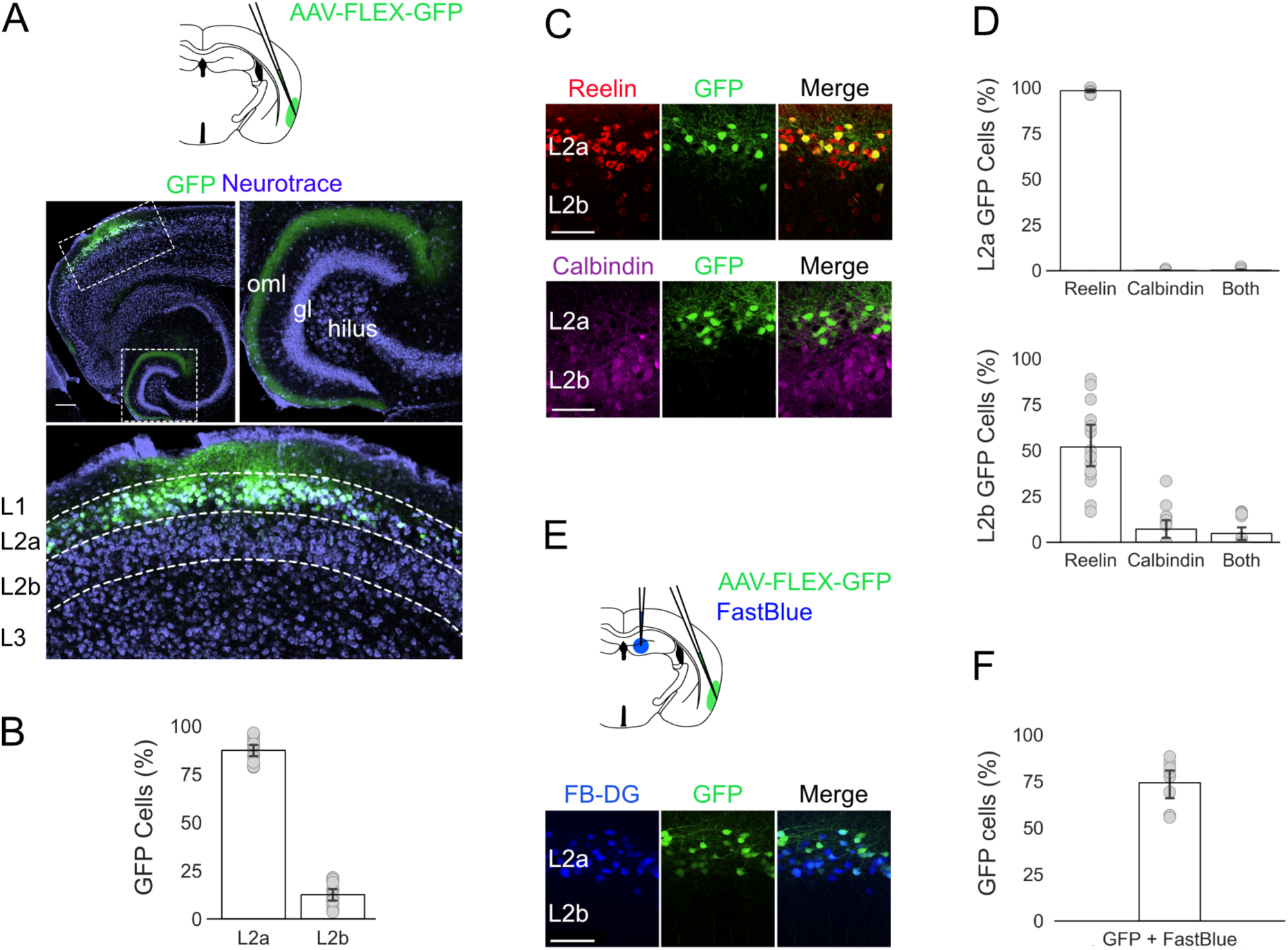
Sim1:Cre mice provide genetic access to reelin cells in L2 of LEC which project to the dentate gyrus. A) Schematic for targeting superficial LEC with AAV-FLEX-GFP and images of a horizontal brain section from a Sim1:Cre mouse showing GFP expression (green) counterstained with Neurotrace (violet). The dashed boxes in the upper left image indicate regions shown in the insets. Axonal labelling is found in the outer molecular layer of the dentate gyrus (upper right) and L2 of LEC (lower). Abbreviations: gl, granule cell layer; oml, outer molecular layer. The scale bar is 250 µm. B) Percentage of neurons that expressed the reporter gene that were in L2a (left, n = 4 mice, 87.4 ± 1.6%, 1282/1490 cells, 14 sections) and L2b (right, 12.6 ± 1.6%, 207/1490 cells) of the LEC. Grey dots indicate percentage values calculated for each section of tissue. Error bars represent SEM. C) Examples of AAV-FLEX-GFP labelled cells which are positive for reelin (red, L2a: 98.3 ± 0.4%, 1260/1282 cells; L2b: 52.0 ± 5.7%, 97/207 cells), but not calbindin (purple, L2a: 0.1 ± 0.1%, 1/1282 cells; L2b: 7.1 ± 2.8%, 17/207 cells). Scale bars represent 100 μm. D) Proportion of neurons expressing the reporter gene (GFP) that were labelled by staining with antibodies against reelin and calbindin. Grey dots indicate percentage values calculated for each section of tissue. Error bars represent SEM. E) Cells labelled by injection of retrograde tracer (blue) overlap with cells labelled by injections of AAV-FLEX-GFP. Scale bar represents 100 μm. Schematic shows the injection strategy used to target superficial LEC with AAV-FLEX-GFP (green) and the dorsal dentate gyrus with a retrograde tracer (Fastblue, blue). F) Proportion of neurons expressing the reporter gene (GFP) that were back-labelled by the injection of retrograde tracer into the dentate gyrus (74.3 ± 4.4%, 331/441 cells, 8 sections). Grey dots indicate percentage values calculated for each tissue section. Error bar represents SEM.

L2 of LEC contains several morphologically and electrophysiologically distinct subtypes of neuron, including pyramidal, multi-form, and fan cells (Tahlvidari & Alonso, 2005; Canto & Witter, 2012; Leitner et al. 2016). To establish whether the population of cells labelled in Sim1:Cre mice mapped onto a specific subtype, we injected AAV encoding a Cre-dependent fluorescent reporter (AAV-hSyn-DIO-hM4D(Gi)-mCherry) into the superficial LEC of Sim1:Cre mice (n = 5) and performed patch-clamp recordings in brain slices from labelled cells in L2a of LEC (n = 15 cells). The electrophysiological properties of the labelled cells were consistent with those previously described for fan cells (Figure 2A; Supplemental Table 2.1), including a relatively depolarized resting membrane potential (−68.5 ± 1.4 mV), high input resistance (130.8 ± 11.9 mΩ), slow time constant (24.2 ± 1.7 ms) and a ‘sag’ membrane potential response to injection of hyperpolarizing current (0.85 ± 0.1). Reconstruction of a subset of these cells (n = 12) revealed that their dendritic architecture was similar to previous descriptions of fan cells, with a polygonal soma and ‘fan-like’ arrangement of primary dendrites. To test whether these cells provide synaptic output to the hippocampus, we injected AAV encoding Cre-dependent channelrhodopsin-2 (ChR2; AAV-EF1a-DIO-hChR2(H134R)-EYFP) into the superficial LEC of Sim1:Cre mice (n = 5) to enable their optical activation (Figure 2B). We then recorded light-evoked glutamatergic synaptic potentials in downstream granule cells in the dentate gyrus (Figure 2C; n = 6 cells), confirming that the labelled cells mediate perforant path inputs from the LEC to the dentate gyrus.

**Figure 2:**
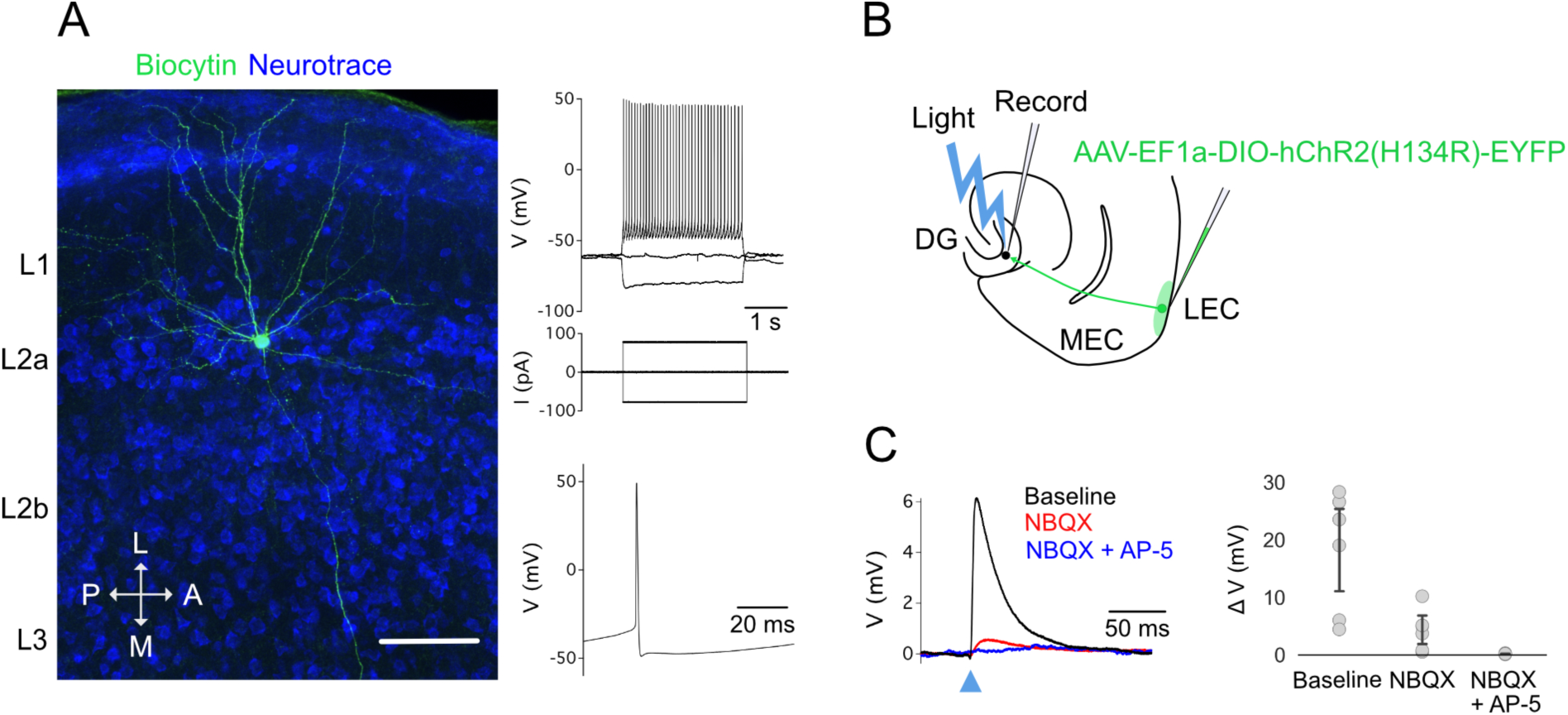
Neurons labelled by Sim1:Cre mice in layer 2 of LEC are fan cells. A) Representative example of the morphology and electrophysiology of a fan cell which expressed mCherry after injection of AAV-hSyn-DIO-hM4D(Gi)-mCherry into the superficial LEC of a Sim1:Cre mouse. (Left) The dendrites and axons of the cell are filled with biocytin (green) and neurons are counterstained with Neurotrace (blue). Note the axon leading towards the dentate gyrus. Scale bar represents 100 μm. (Right) Membrane potential response to the injection of negative and positive current steps (top) and example action potential (bottom). B) Schematic of recording experiment to confirm that fan cells labelled in Sim1:Cre mice project to the dentate gyrus. AAV-EF1a-DIO-ChR2(H134R)-EYFP was injected into the superficial LEC of Sim1:Cre mice (n = 5). Synaptic output from fan cells in layer 2 of LEC was evaluated by recording light evoked responses of granule cells in the dentate gyrus. C) Membrane potential response of a dentate gyrus granule cell after optogenetic activation of fan cells in layer 2 of LEC expressing ChR2. (Left) The peak response was abolished by application of ionotropic glutamate receptor antagonists NBQX (red) and AP-5 (blue). (Right) Quantification of the light evoked membrane potential response of dentate gyrus granule cells (n = 6) after application of NBQX and AP-5. Each grey circle represents an individual neuron. Error bars are SEM.

Together, these experiments demonstrate that Sim1:Cre mice can be used to obtain genetic access to fan cells in L2 of the LEC. Our analyses confirm that fan cells express reelin and project to the dentate gyrus of the hippocampus.

### Fan cells in L2 of LEC are critical for episodic-like memory

To test whether fan cells are required for associative recognition memory, we blocked their output using targeted injections of AAV encoding a Cre-dependent tetanus toxin light chain (TeLC) (Figure 3A; AAV-FLEX-TeLC-GFP), an approach we previously validated for inactivation of stellate cells in L2 of the MEC (Tennant et al. 2018). We compared Sim1:Cre mice injected with AAV-FLEX-TeLC-GFP (n = 21) with corresponding control mice injected with AAV-FLEX-GFP (n = 17).

**Figure 3:**
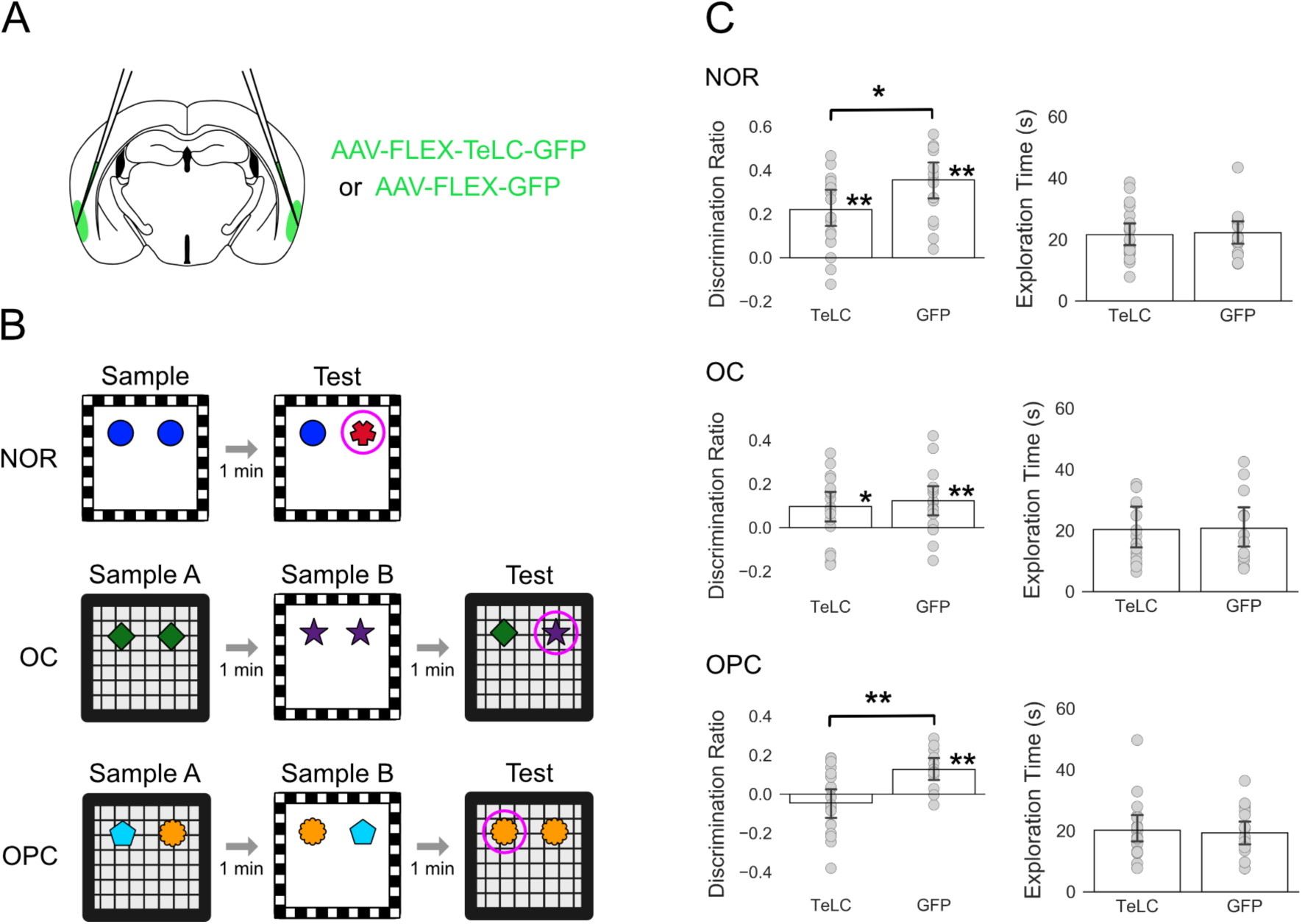
Suppression of fan cells in L2 of LEC impairs performance on the object-place-context task. A) Schematic of injection strategy used to bilaterally target superficial LEC with AAV-FLEX-TeLC-GFP or AAV-FLEX-GFP. B) Schematic of object recognition tasks. Tasks included novel object recognition (NOR, top), object-context (OC, middle) and object-place context (OPC, bottom). Purple circles highlight the novel object or configuration in the test trial. C) Average discrimination ratios (D’) and exploration times for each task. Discrimination ratios reflect the proportion of time spent exploring the novel object or configuration (Ennaceur et al. 1980). (Left) Bars indicate average discrimination ratio for each group. One-sample *t-*tests against a value of 0 revealed that mice in both groups performed significantly above chance on the NOR task (top, TeLC: *t*(18) = 5.001, *P* < 0.001; GFP control: *t*(15) = 7.922, *P* < 0.001) and the OC task (middle, TeLC: *t*(18) = 2.742, *P* = 0.013, GFP control: *t*(15) = 3.340, *P =* 0.004). In contrast, mice in the TeLC group showed no exploratory preference for the novel configuration in the OPC task (bottom, *t*(18) = −1.183, *P* = 0.252), whereas mice in the control group explored the novel configuration more than predicted by chance (*t*(11) = 4.274, *P* = 0.001). Univariate ANOVAs revealed a significant difference in performance between TeLC mice and control mice for the NOR (*P* = 0.041) and OPC tasks (*P* = 0.003), but not the OC task (*P* = 0.603). Asterisks indicate a significant difference between groups or performance that is significantly different from chance (* = *P* < 0.05, ** = *P* < 0.01). (Right) Bars indicate average exploration time in seconds during the test trials for each group. Each grey dot represents the data for a single animal. Error bars represent SEM.

We assessed performance of the mice on a series of object-based memory tasks (Figure 3B), which model the integration of episodic memory components in rodents (Eacott & Norman, 2004). In the sample phase of each task, animals encountered two objects in different spatial locations within an environmental context that has distinct visual and tactile features. After a short interval, the animal was presented with a familiar object and an object which was novel or in a novel configuration of location and/or context.

Mice expressing TeLC in fan cells were able to distinguish novel objects and object-context configurations. Both TeLC and control mice explored novel objects more than expected by chance, although discrimination ratios were lower for the TeLC group (Figure 3C; *F*_(1,33)_= 4.542, *P*= 0.041, η^2^ = 0.121). Novel object-context configurations were also explored more than expected by chance by the TeLC and control mice, with no difference between the two groups (*F*_(1,33)_= 0.275, *P* = 0.603). These data suggest that fan cells are not required for recognition of novel objects, and unlike the LEC as a whole, are not required for recognition of novel object-context associations.

When we investigated discrimination of more complex associations of object, place, and context, we found that mice in the TeLC group showed no exploratory preference for the novel configuration, whereas mice in the control group explored the novel configuration more than predicted by chance. Consistent with this, there was a significant difference in performance between the two groups (*F*_(1,29)_= 10.319, *P* = 0.003, η^2^ = 0.262). The amount of time spent exploring the objects was similar across groups for all tasks, indicating that differences between groups are not explained by reduced interest in the objects.

If deficits in task performance result from expression of TeLC then performance should correlate with the extent to which AAV-FLEX-TeLC-GFP infected fan cells across the LEC. To test this, we quantified for each animal the percentage area of L2 of LEC with virus expression (Figure 4A-B). We found that the pattern of GFP labelling in the older mice used for behavioural experiments was similar to that of younger mice used for electrophysiological and anatomical experiments (cf. Figures 1A and 4A). When we compared the percentage area of virus expression with object-place-context discrimination we found a negative correlation for the TeLC group (Figure 4C; *R* = −0.483, n = 18, *P* = 0.042), but not for the GFP group (*R* = 0.356, n = 9, *P* = 0.347). We did not find any correlation between viral expression and performance for the other recognition tasks (Supplemental Figure 4.1).

**Figure 4:**
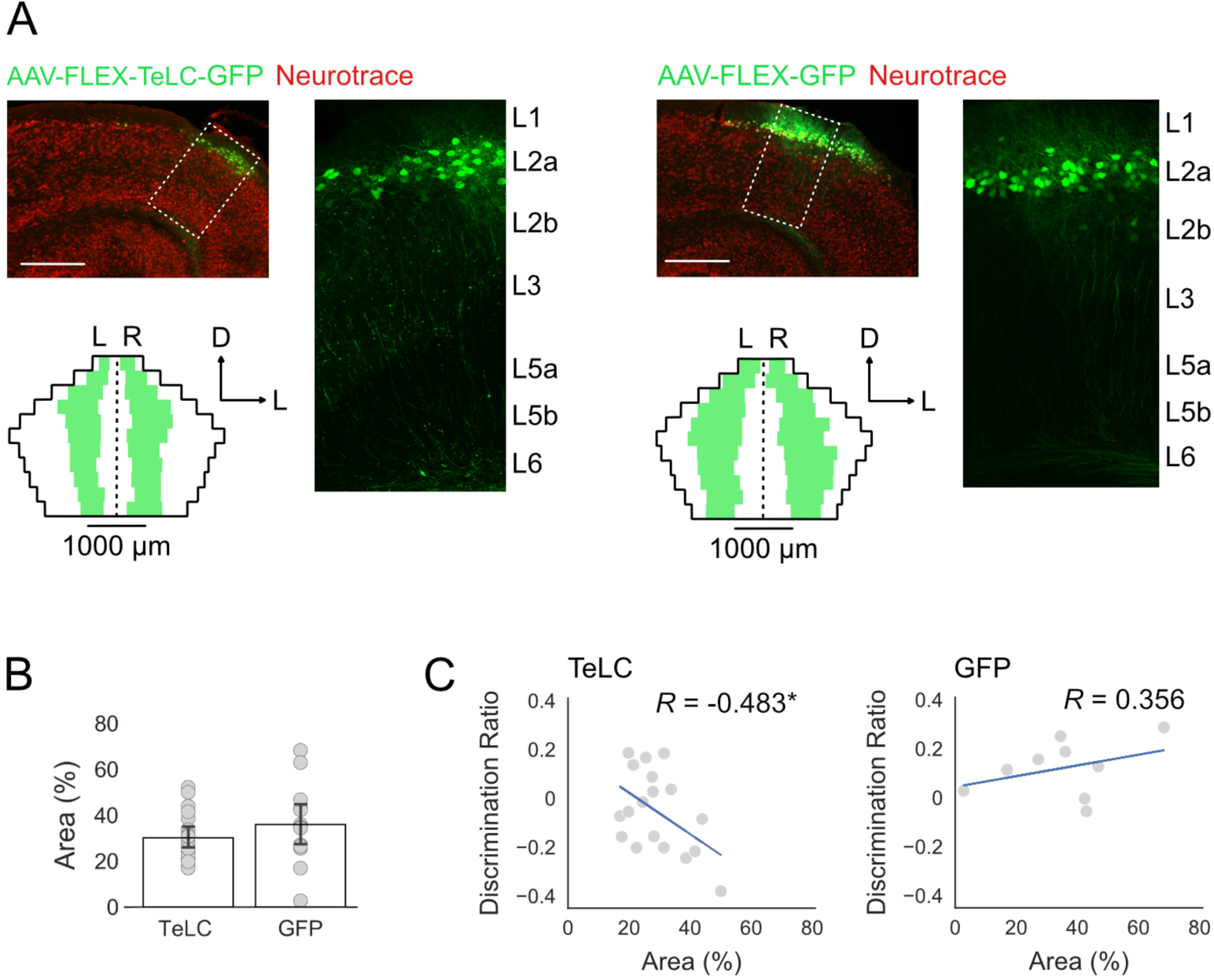
Quantification of virus expression in mice injected with AAV-FLEX-TeLC-GFP or AAV-FLEX-GFP. A) Horizontal brain sections showing expression of AAV-FLEX-TELC-GFP (left) or AAV-FLEX-GFP (right) in a Sim1:Cre mouse (upper left). Reporter gene expression (GFP, green) is shown against neurons counterstained with Neurotrace (red). Scale bar represents 250 µm. The dashed boxes indicate the regions of LEC which are shown in the insets (right). Unfolded representations of LEC L2a (bottom) are overlaid with the average location and spread of virus expression for each group. Average distances between regions of virus expression and the borders of LEC (white) and average lengths of virus expression (green) were calculated by averaging measurements across all animals for sections at each dorsoventral coordinate of LEC (D = dorsal, L = lateral). Left and right hemispheres are indicated by L and R, and black dotted line indicates separation between hemispheres. B) Bars indicate average percentage area of L2 of LEC which contained neurons that expressed the reporter gene after injection of AAV-FLEX-TELC-GFP or AAV-FLEX-GFP. Each grey dot represents the data for a single animal. Error bars are SEM. C) Relationship between virus expression and performance for the OPC task. Scatterplots of the percentage area of virus expression in LEC L2 plotted against the discrimination ratios for the OPC task. Each grey dot represents data for a single animal. Pearson’s correlation coefficients (*R*) were calculated to determine the strength of the relationship between discrimination ratios and virus expression. There was a significant negative correlation between discrimination ratios and virus expression in the TeLC group (*P* = 0.042), but not the GFP group (*P* = 0.347). Asterisks indicate a significant correlation between discrimination ratios and virus expression (* = *P* < 0.05). Each plot is overlaid with the line of best fit (blue) that was calculated using the least squares method of linear regression.

## Discussion

Our data provide evidence that fan cells in L2 of LEC are critical for episodic-like memory. This is in contrast to complete lesions of the LEC, which cause a general associative memory impairment (Wilson et al. 2013a; 2013b). Our data are consistent with a role of the LEC in encoding complex information about objects in the environment (Deshmukh et al. 2011; Tsao et al. 2013; 2018; Wang et al. 2018), and suggest that this information may contribute to the formation of object-place-context associations via projections from fan cells to the hippocampus. Impaired episodic-like memory but spared recognition of simpler associations following lesions of the hippocampus are consistent with this possibility (Langston & Wood, 2010).

Functional specialisation within LEC circuits has implications for normal circuit computations and disorders. Our results suggest a model in which L2a generates outputs specifically required for relatively complex object-place-context associations, whereas neurons in other layers maybe sufficient for formation of simpler associations. Pathology of the superficial LEC has been associated with early cognitive deficits in AD and in age-related cognitive decline (Gomez-Isla et al. 1996; Stranahan et al. 2011a; 2011b; Khan et al. 2014; Kobro-Flatmoen et al. 2016). By showing that episodic-like memory requires output from fan cells, our results suggest that pathology specific to L2a of LEC is sufficient to account for associative memory impairments in early stages of AD.

## Materials and Methods

### Animals

The Sim1:Cre line, which expresses Cre under the control of the Single minded homolog-1 (Sim1) promoter, was generated by GenSat and obtained from MMRRC (strain name: Tg(Sim1cre)KH21Gsat/Mmucd). Sim1:Cre mice were bred to be heterozygous for the Cre transgene by crossing a male Sim1:Cre mouse carrying the transgene with female C57BL6/J mice. Anatomical and electrophysiological experiments used 5-11 week old mice and the behavioural experiment used 3-8 month old mice. All mice were housed in groups in diurnal light conditions (12-hr light/dark cycle) and had *ad libitum* access to food and water. All experiments and surgery were conducted under project licenses administered by the UK Home Office and in accordance with national (Animal [Scientific Procedures] Act, 1986) and international (European Communities Council Directive of 24 November 1986 (86/609/EEC) legislation governing the maintenance of laboratory animals and their use in scientific research.

### Viruses

We used the following adeno-associated viruses: AAV-1/2-FLEX-GFP and AAV-1/2-FLEX-TeLC-GFP (generated in house following protocols described by Murray et al. 2011), AAV2-hSyn-DIO-hM4D(Gi)-mCherry (UNC Vector Core, Chapel Hill, North Carolina, Lot #: AV44362) and AAV2-EF1a-DIO-hChR2(H134R)-EYFP (UNC Vector Core, Chapel Hill, North Carolina, Lot #: AV43780).

### Stereotaxic Injection of Tracers and Viruses

Mice were anaesthetised with Isoflurane in an induction chamber before being transferred to a stereotaxic frame. Mice were administered an analgesic subcutaneously, and an incision was made to expose the skull. For retrograde labelling of neurons which project to the hippocampus, Fast Blue (Polysciences, Hirschberg an der Bergstraße, Germany) was injected unilaterally into the dorsal dentate gyrus. A craniotomy was made at 2.8 mm posterior to bregma and 1.8 mm lateral to the midline. A glass pipette was lowered vertically into the brain to a depth 1.7 mm from the surface, and 50-100 nl of tracer diluted at 0.5% w/v in dH_2_O was injected. For injection of virus into the lateral entorhinal cortex, a craniotomy was made adjacent to the intersection of the lamboid suture and the ridge of the parietal bone, which was approximately 3.8 mm posterior to bregma and 4.0 mm lateral to the midline. From these coordinates, the craniotomy was extended 0.8 mm rostrally. At the original coordinates, a glass pipette was lowered from the surface of the brain at a 10°-12°angle until a slight bend in the pipette indicated contact with the dura. The pipette was retracted 0.2 mm and 150-500 nl of virus was injected. This protocol was repeated at a site 0.2 mm rostral to this site. To target ventral lateral entorhinal cortex, the angle of the pipette was adjusted to 6° - 9° and a third injection was delivered at the rostral injection site. The angle was adjusted within each reported range based on the proximity of the craniotomy to the ridge of the parietal bone. This approach minimised the likelihood of spread of virus into adjacent cortical structures. For all injections, the pipette was slowly retracted after a stationary period of four minutes. Mice were administered an oral analgesic prepared in flavoured jelly after surgery.

### Immunohistochemistry

For tracer injections, animals were sacrificed 1-2 weeks post-surgery. For anatomical and electrophysiological experiments animals were sacrificed 2-4 weeks post-surgery. Mice were administered a lethal dose of sodium pentobarbital and transcardially perfused with cold phosphate buffered saline (PBS) followed by cold paraformaldehyde (PFA, 4%). Brains were extracted and fixed for a minimum of 24 hours in PFA at 4°C, washed with PBS, and transferred to a 30% sucrose solution prepared in PBS for 48 hours at 4°C. Brains were sectioned horizontally at 60 μm on a freezing microtome. For staining against reelin and calbindin, slices were blocked for 2-3 hours in 5% Normal Goat Serum (NGS) prepared in 0.3% PBS-T (Triton). Slices were then transferred to a primary antibody solution prepared in 1% NGS in 0.3% PBS-T for 24 hours. Primary antibodies were mouse anti-reelin (MBL, 1:200, Catalogue #: D351-3) and rabbit anti-calbindin D-28K (SWANT, 1:2500, Catalog #: CB-38). Slices were washed with 0.3% PBS-T 3x for 20 minutes before being transferred to a secondary antibody solution prepared in 0.3% PBS-T for 24 hours. Secondary antibodies were AlexaFluor® Goat Anti-Mouse 488 (Catalog #: A-10680 and 546 (Catalog #: A-11003) and AlexaFluor® Goat Anti-Rabbit 555 (Catalog #: A27039) and 647 (Catalog #: A27040; Invitrogen, all used at 1:500). Neurotrace 640/660 (Catalog #: N21483; Invitrogen, 1:800) was included in the secondary antibody solution as a counterstain. Slices were washed with 0.3% PBS-T 3x for 20 minutes and then mounted and cover-slipped with Mowiol.

### Quantification of Immunohistochemistry

All images were acquired using a Nikon A1 confocal microscope and NIS elements software. For quantification of cells immunolabelled for reelin or calbindin, or labelled by retrograde tracer or fluorescent reporter, 20x z-stacks were acquired of regions of interest (ROI) at 1–2 μm steps. ROIs were regions of L2 of various sizes within the borders of LEC, which were determined by referencing an atlas of the mouse brain (Paxinos & Franklin, 2007). All neurons within each ROI were counted manually. Fractions of labelled cells were determined by calculating for each ROI the number of antibody labelled cells divided by the total number of neurotrace labelled cells. Proportions were averaged across all ROIs from all mice to yield an overall percentage of labelled cells.

### Slice Electrophysiology

Horizontal brain slices were prepared from 5-11 week old Sim1:Cre mice 2-4 weeks after injection of AAV-hSyn-DIO-hM4D(Gi)-mCherry or AAV-EF1a-DIO-hChR2(H134R)-EYFP into superficial LEC. Mice were sacrificed by cervical dislocation and decapitated. The brains were rapidly removed and submerged in cold cutting artificial cerebrospinal fluid (ACSF) at 4-8°C. The cutting ACSF was comprised of the following (in mM): NaCl 86, NaH_2_PO4 1.2, KCl 2.5, NaHCO_3_25, Glucose 25, Sucrose 50, CACl_2_ 0.5, MgCI_2_ 7. The dorsal surface of the brain was glued to a block submerged in cold cutting ACSF and horizontal slices were cut using a vibratome at a thickness of 300-400 μm. Slices were transferred to standard ACSF at 37°C for 15 minutes, then incubated at room temperature for a minimum of 45 minutes. Standard ACSF was comprised of the following (in mM): NaCl 124, NaH_2_PO4 1.2, KCI 2.5, NaHCO_3_ 25, Glucose 25, CaCI_2_ 2, MgCI_2_ 1. For recordings, slices were transferred to a submerged chamber and maintained in standard ACSF at 35-37°C. Labelled neurons in LEC were identified by their expression of the relevant fluorophore. Neurons were patched in the granule cell layer of the dentate gyrus in regions where there was axonal expression of the virus visible in the outer molecular layer. Whole cell patch-clamp recordings were made using borosilicate electrodes with a resistance of 3-6 MΩ for cells in LEC and 6-10 MΩ for cells in dentate gyrus. Electrodes were filled with an intracellular solution comprised of the following (in mM): K gluconate 130, KCI 10, Hepes 10, MgCl_2_2, EGTA 0.1, Na_2_ATP 4, Na_2_GTP 0.3, phosphocreatine 10, and 0.5% biocytin (w/v). Recordings were made in current-clamp mode from cells with series resistance ≤ 50 MΩ with appropriate bridge-balance and pipette capacitance neutralisations applied.

### Recording Protocols

A series of protocols were used to characterise the electrophysiological properties of each cell recorded in LEC or the dentate gyrus, as described previously (Nolan et al. 2007; Garden et al. 2008). Sub-threshold membrane properties were measured by examining membrane potential responses to injection of current in hyperpolarizing and depolarizing steps (−160 to 160 pA in 80 pA increments, or −80 to 80 pA in 40 pA increments, each 3 s duration), and to injection of an oscillatory current with a linearly varying frequency (ZAP protocol). Supra-threshold properties were estimated from responses to depolarizing current ramp applied to the cell in a constantly increasing manner to induce action potentials (50 pA/s, 3s). Responses to optogenetic activation of inputs to dentate granule cells were evaluated by stimulation with 470 nm wavelength light for 3 ms at 22.4 mW/mm^2^. After recording a baseline response to light stimulation, ionotropic glutamate receptor antagonists for AMPA (NBQX, 10 µM) and NMDA receptors (AP-5, 50 µM) were bath applied, and the response to light stimulation was re-evaluated using the same stimulation protocols. Upon completion of investigatory protocols, diffusion of biocytin into the cell was encouraged by injecting large depolarizing currents into the cell (15 x 4 nA, 100 ms steps, 1Hz; Jiang et al. 2015). Each cell was left stationary with the electrode attached for a maximum of one hour before being transferred to PFA (4%) and stored at 4°C for at least 24 hours before histological processing.

### Analysis of Electrophysiological Data

Electrophysiological data were analysed with AxoGraph (axographx.com), IGOR Pro (Wavemetrics, USA) using Neuromatic (http://www.neuromatic.thinkrandom.com), and customised MATLAB scripts. Input resistance, time constant and time-dependent inward rectification (‘sag’) were measured from the membrane potential response to injections of hyperpolarizing current (−80 or −40 pA). Input resistance (MΩ) was estimated by dividing the steady-state voltage change from the resting membrane potential by the amplitude of the injected current, time constant (ms) was measured as the time taken for the change in voltage to reach a level 37% above its maximal decrease, and ‘sag’ was measured as the ratio between the maximum decrease in voltage and the steady-state decrease in voltage. Rheobase (pA) was measured as the minimum amplitude of depolarizing current which elicited an action potential response from the cell. Action potential duration (ms) was measured from the action potential threshold (mV), which was defined as the point at which the first derivative of the membrane potential voltage that exceeded 1 mV/1 ms. Action potential amplitude (mV) was measured as the change in voltage between the action potential threshold and peak. To determine the resonant frequency of the cell, membrane impedance was first calculated by dividing the Fourier transform of the membrane voltage response by the Fourier transform of the input current from the ZAP protocol, which was then converted into magnitude and phase components (see Nolan et al. 2007). The resonant frequency was defined as the input frequency which corresponded to the peak impedance magnitude. To quantify the response of granule cells to optical stimulation, the change in amplitude was measured between the resting membrane potential and peak response during stimulation.

### Neuron Reconstruction

To reveal the morphology of biocytin filled neurons, fixed slices were washed with PBS 4 times for 10 minutes and transferred to a solution containing AlexaFluor® Streptavidin 488 (Invitrogen, 1:1000 or 1:500, Catalog **#:** S32354) and Neurotrace 640/660 (Invitrogen, 1:1000 or 1:500) in 0.3% PBS-T for 24 hours. Slices were washed with PBS-T 4 times for 20 mins and then mounted and cover-slipped with Mowiol. Z-stacks were acquired of filled cells at 1-2 μm steps using a 20x objective on either Nikon A1 or Zeiss LSM800 confocal microscopes. The morphology of cells recorded in LEC were confirmed by visual comparison of the shape of the soma and arrangement of dendrites to published morphological descriptions (Tahvildari & Alonso, 2005; Canto & Witter, 2012; Leitner et al. 2016).

### Behavioural Apparatus

The test environment was a rectangular wooden box (length 32 cm, width 25.5 cm, height 22 cm) that could be configured to provide two distinct contexts. Context A had black and white vertically striped walls with a smooth white floor. Context B had grey walls with a wire-mesh floor. The environment was lit by two lamps positioned equidistantly from the box. The wall and floor of the environment was cleaned with veterinary disinfectant before each trial. To secure objects in place within the environment, square sections of fastening tape (Dual Lock, 3M) were positioned in each of the upper left and right quadrants of the box floor. Objects were household items of approximately the same size as a mouse and varying in colour, shape, and texture. Objects were matched for similarity in size and complexity when paired for testing. To avoid the use of odour cues, new identical copies of each object were used for each trial, and objects were cleaned with disinfectant after each trial.

### Design of Behavioural Experiments

Behavioural experiments were carried out in two cohorts of mice. In the main manuscript data from the cohorts are pooled. To control for order and cage effects, each cage contained a mixture of mice from the experimental and control group. The experimenter was blind to the manipulation (delivery of either AAV-FLEX-TeLC-GFP or AAV-FLEX-GFP) during all behavioural experiments and their initial analyses.

### Habituation

In the week before surgery, the experimenter handled each animal every day for 10 minutes in the testing room. Habituation to the test environment commenced one week after surgery and lasted for 5 consecutive days. On day 1, the mice explored each context for 10 minutes with their cage mates. On days 2-5 the mice explored each context for 10 minutes individually. On days 4-5, objects were introduced in the upper left and right quadrants of the test environment. Objects used during habituation were not re-used during testing.

### Behavioural Tasks

Testing commenced in the week after habituation and occurred in three stages: Novel object recognition (NOR), novel object-context (OC) recognition, and novel object-place-context (OPC) recognition. Novel object-place recognition was also included as a stage in these experiments, but neither experimental group of mice performed above chance level in this task therefore we excluded it from further analysis. Each stage lasted for 4 days. For all sample and test trials, the animal was allowed to explore the environment freely for 3 mins. Between trials, mice were removed to a holding cage for approximately 1 minute while the test environment was configured for the subsequent trial. For each task, the object that was novel at test, the context, and the quadrants where the novel object or configuration occurred were counterbalanced across animals and experimental conditions. For OC and OPC tasks, the presentation order of context A and B in the sample phase, the context used at test, and the context in which each object was presented during the sample phase were also counterbalanced across animals and conditions. The stages are described here in the order in which they occurred:

1. NOR Task: In the sample trial, mice were presented with two copies of a novel object in one of the contexts (striped or grey walls). In the test trial, mice were presented with a new copy of the object from the sample trial (now familiar) and a novel object in the same context as the sample trial.
2. OC task: In the first sample trial, mice were presented with two copies of a novel object in context A. In a second sample trial, mice were presented with two copies of a different novel object in context B. In the test trial, mice were presented with a single copy of each object within one of the contexts (A or B). At test, both objects are familiar and have been encountered at both locations, but one object has previously been encountered in the test context (familiar OC configuration), whereas the other one has not been encountered in the test context (novel OC configuration).
3. OPC task: In the first sample trial, mice were presented with two different novel objects in context A. In a second sample trial, mice were presented with the same objects in opposite locations in context B. In the test trial, mice were presented with two copies of one of the objects within one of the contexts (A or B). At test, one copy of the object is in a novel location for the test context (novel OPC configuration), whereas the other copy is in a familiar location for the test context (familiar OPC configuration).

### Histology and Quantification of Virus Expression

Mice were transcardially perfused as described previously. Slices were washed in 0.3% PBS-T 3 times for 20 minutes before being transferred to a solution containing Neurotrace 640/660 (Invitrogen, 1:800) prepared in 0.3% PBS-T for 2-3 hours at room temperature. Slices were washed with 0.3% PBS-T 3 times for 20 minutes before being mounted and cover-slipped with Mowiol. Mounted sections were stored at 4°C. All images were acquired using a fluorescent microscope (Zeiss ApoTome) and ZenPro software. To confirm the location and extent of virus expression in each animal, 1:4 sections which contained LEC were imaged at a 10x magnification. Coordinates for each section were determined by referencing an atlas of the mouse brain (Franklin & Paxinos, 2007). Unfolded representations of LEC L2a were generated by adapting procedures previously used to quantify lesions of the entorhinal cortex (Insausti et al. 1997; Steffenach et al. 2005). The anterior and posterior borders of the LEC were determined by the bifurcation of L2, which is absent in adjacent cortical structures. A subset of brains (n = 1 TeLC, 4 GFP) suffered mechanical damage to LEC L2 in > 30% of sections that contained LEC. Removal of these mice from the dataset did not change the interpretation of statistical comparison between groups, therefore they were included in the analysis of behaviour but excluded from analyses which address the relationship between virus expression and behaviour. The length of L2a was measured in ImageJ (https://fiji.sc) using a built-in tool calibrated to the scale of the image. For sections with virus expression, three measurements were extracted: the distance of the region of expression from the anterior LEC border, the length of the region of expression, and the distance of the region of expression from the posterior LEC border. The region of expression was defined as the length of L2a between the most anterior infected neuron and the most posterior infected neuron. These measurements were used to calculate the proportion of LEC L2a in which the virus was expressed for each animal, and averaged across all animals for sections with virus expression across the dorsoventral axis to obtain the mean values used to generate the unfolded representations shown in Figure 4A.

Adjacent structures were examined for unintended expression of virus in the TeLC group. There was labelling in LEC L5a in a subset of mice (n = 11). In most cases (n = 8), this was negligible, summating to <10 cells across all sections. In other cases (n = 3), this was approximated to be <5% of the area of LEC L5. Further, there was spread of virus to the region of MEC L2 directly adjacent to LEC in a subset of mice (n = 8), but this was approximated to be <5% of the area of MEC L2 in all cases. There was no significant difference between performance of mice with L5 expression or MEC expression and other mice in the TeLC group on any task. To determine the density of virus expression in each slice, each region of expression was imaged at a 20x magnification. For each region, the number of neurons expressing the reporter gene (GFP) and the number of counterstained neurons were quantified manually. From these values, the percentage of infected neurons within each region of expression was calculated as the number GFP of labelled cells divided by the number of Neurotrace labelled cells in the region. These values were averaged across all quantified sections to produce a single density value for each mouse.

### Analysis of Behavioural Data

All trials were recorded by a camera positioned above the test environment. Footage was scored offline by an experimenter who was blind to experimental condition. To ensure reliability of the scores a random sample of 50% of the trials were rescored by a second experimenter who was also blind to condition. Reliability between scorers was good with an intraclass correlation coefficient of 0.878 (2-way mixed model). For each trial, the amount of time spent exploring each object was measured. Exploration was defined as periods where the animal’s nose is within 2 cm of the object and oriented towards the object, but the animal was not interacting with the object (eg. biting) or rearing against the object to look out of the test environment. To determine whether the animal had an exploratory preference for the novel object or configuration, a discrimination ratio was calculated for each test trial (Ennaceur & Delacour, 1980). The discrimination ratio is calculated by subtracting the amount of time spent exploring the object in the familiar configuration from the amount of time spent exploring the object in a novel configuration, and then dividing this value by the total exploration time. A positive discrimination ratio indicates an exploratory preference for the object in a novel configuration. For each animal, average discrimination ratios were calculated for each task. A population mean was then calculated for experimental (AAV-FLEX-TeLC-GFP) and control (AAV-FLEX-GFP) groups. Trials where the total exploration time was < 5 seconds during sample or test were excluded. Where ≥ 3 trials of a task met the criteria for exclusion for an animal, data from that animal was removed from the dataset for that task (NOR, n = 3, 1 TeLC and 2 GFP control; OC, n = 2 GFP control, OPC: n = 7, 2 TeLC and 5 GFP control). Behavioral data are provided in the Supplemental Data document.

### Statistical Analysis of Behavioural Data

All statistical analyses were conducted using SPSS (IBM, version 24). To determine whether there was an effect of experimental group, univariate ANOVAs with group (experimental and control) as a between-subjects factor were used for each task. To determine whether the average discrimination ratios for each group were different from chance, one-sample *t*-tests were conducted against a value of 0 for each task. To determine whether there was a relationship between the extent of virus expression and behaviour, Pearson’s product-moment correlation coefficients were calculated with percentage area of virus expression as a variable against the average discrimination ratio of each task for each animal. Lines of best fit were calculated for the dataset using the least squares method of linear regression.

## Author Contributions

Conceptualization and Methodology, B.V., M.F.N., J.A.A., Investigation, B.V., D.L.F.G., V.A., Resources, C.M., Writing - Original Draft, B.V., Writing, Review & Editing, B.V., M.F.N., J.A., Supervision, M.F.N. and J.A., Funding Acquisition, B.V., M.F.N., J.A.

## Funding Acknowledgements

This work was supported by a Carnegie Trust Collaborative Research Grant to J.A. and M.F.N, a Henry Dryerre scholarship from the Royal Society of Edinburgh to B.V., and grants from Wellcome Trust (200855/Z/16/Z) to M.F.N, and BBSRC (BB/M025454/1) to M.F.N.

---

**Supplemental Figure 1.1:**
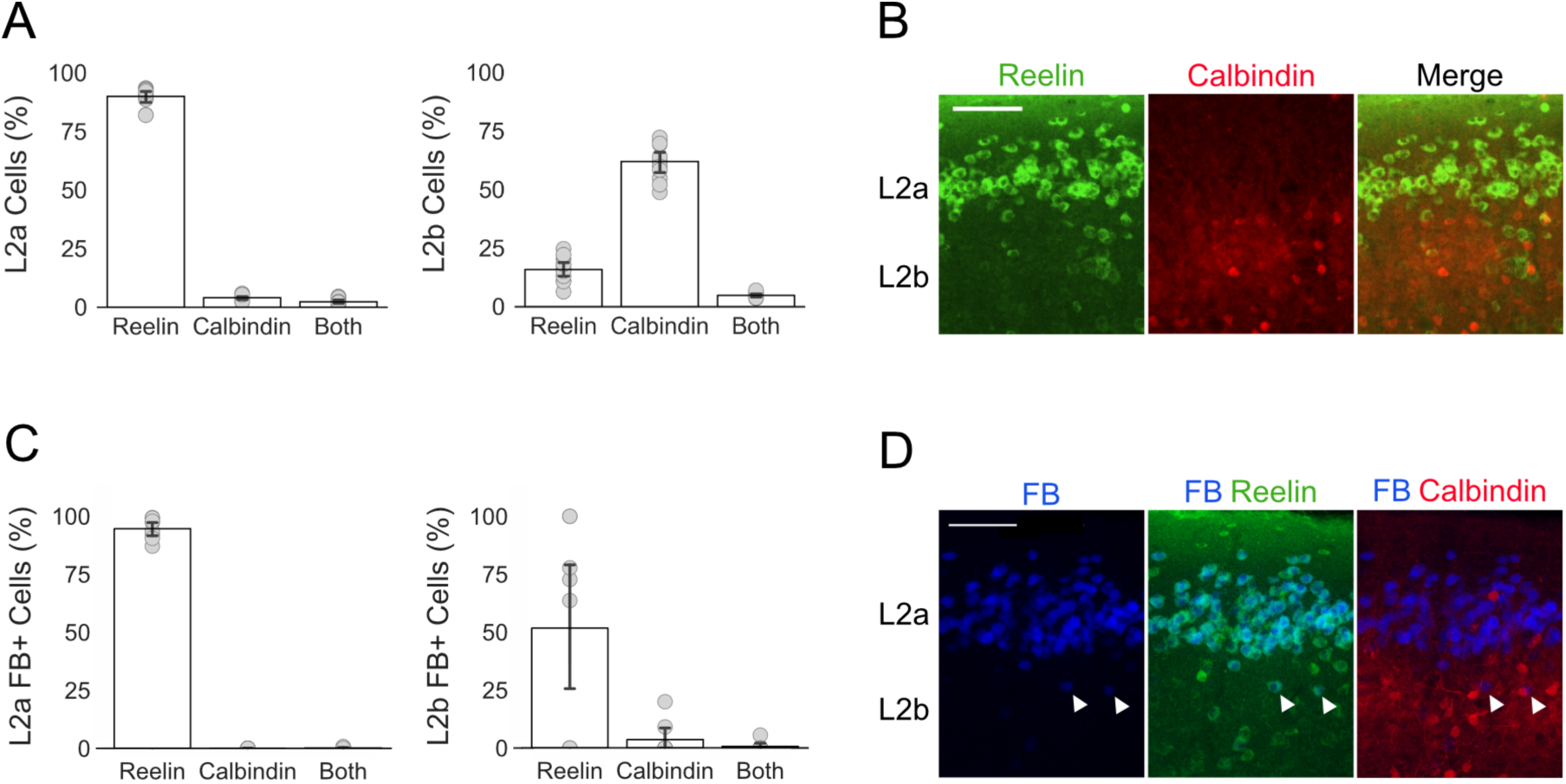
Quantification of reelin, calbindin and retrograde tracer labelling across sub-layers of LEC L2. A) Quantification of cells positive for reelin and calbindin in LEC L2a (left) and L2b (right; n = 4 mice). Note that the majority of cells in LEC L2a and L2b are positive for reelin and calbindin, respectively, as reported by Leitner et al. (2016). Grey dots indicate percentage values calculated for each section of tissue. Error bars represnt SEM. B) Immunolabelling against reelin (green) and calbindin (red) in L2a and L2b of LEC. C) Quantification of cells back-labelled by the retrograde tracer which were positive for reelin and calbindin in LEC L2a (left) and L2b (right; n = 4 mice). Note that the majority of back-labelled cells are positive for reelin in both sub-layers. Grey dots indicate percentage values calculated for each section of tissue. Error bars represent SEM. D) Immunolabelling against reelin (green) and calbindin (red) overlaid with neurons that were back-labelled by the injection of the retrograde tracer FastBlue (FB) into the dentate gyrus (blue). White arrows indicate back-labelled neurons in LEC L2b which are positive for reelin. Scale bars represent 100 µm.

**Supplemental Figure 1.2:**
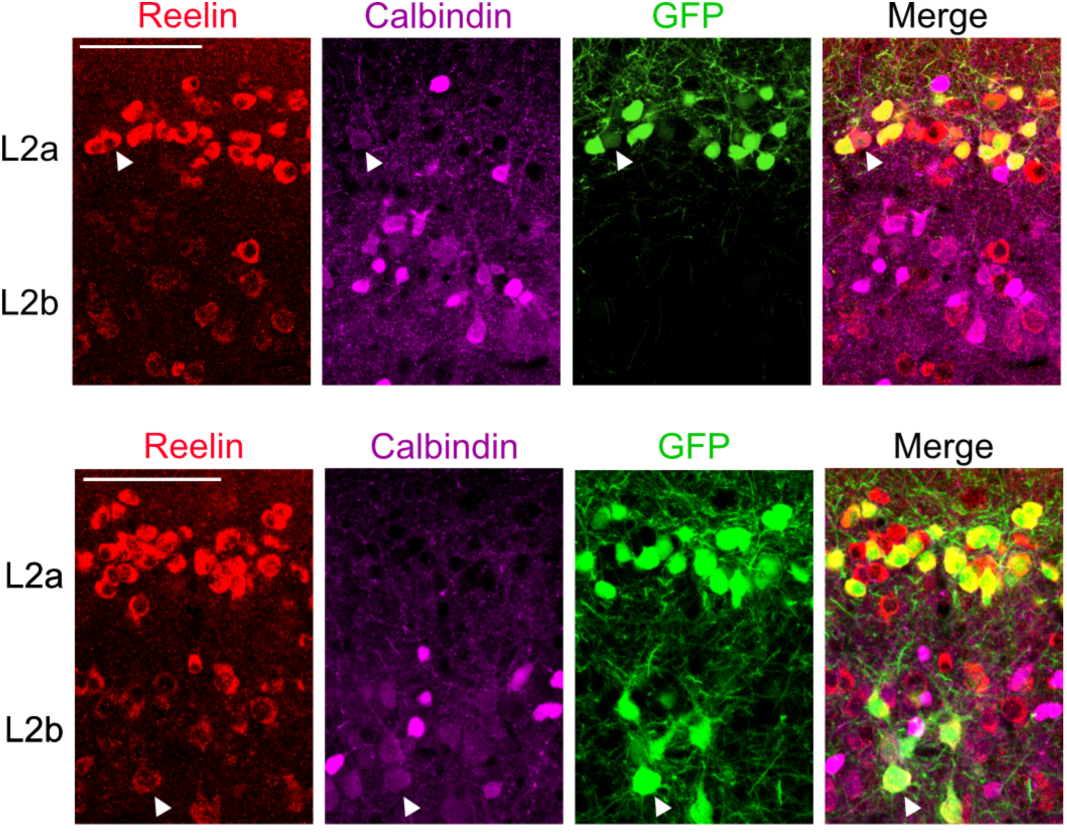
Triple-labelling of neurons in L2 of LEC. Cells triple-labelled by reelin (red), calbindin (purple) and the reporter gene (GFP, green) are indicated by white arrows. A small population of cells was triple-labelled in L2a (top, 0.3 ± 0.2%, 4/1282 cells) and L2b (bottom, 4.6 ± 1.9%, 13/207 cells). Scale bars represent 100 µm.

**Supplemental Table 2.1:** Electrophysiological properties of fan cells. Table contains the population average, standard deviation (SD) and standard error of the mean (SEM) values for the electrophysiological properties of fan cells (n= 15, 5 mice) in LEC L2 which expressed the reporter gene (mCherry) after injection of AAV-hSyn-DIO-hM4D(Gi)-mCherry into the superficial LEC.

| Property | Average | SD | SEM |
| --- | --- | --- | --- |
| Resting membrane potential (mV) | -68.47 | 5.22 | 1.35 |
| Input resistance (mΩ) | 130.47 | 45.97 | 11.87 |
| Time constant (ms) | 24.19 | 6.64 | 1.71 |
| Sag | 0.85 | 0.07 | 0.02 |
| Rheobase (pA) | 126.43 | 34.36 | 8.87 |
| Action potential threshold (mV) | -37.95 | 5.84 | 1.51 |
| Action potential width (ms) | 0.67 | 0.07 | 0.02 |
| Action potential amplitude (mV) | 88.19 | 3.65 | 0.94 |
| Resonance frequency (Hz) | 1.15 | 0.35 | 0.09 |
| Resonance magnitude (mV) | 1.03 | 0.05 | 0.01 |

**Supplemental Figure 4.1:**
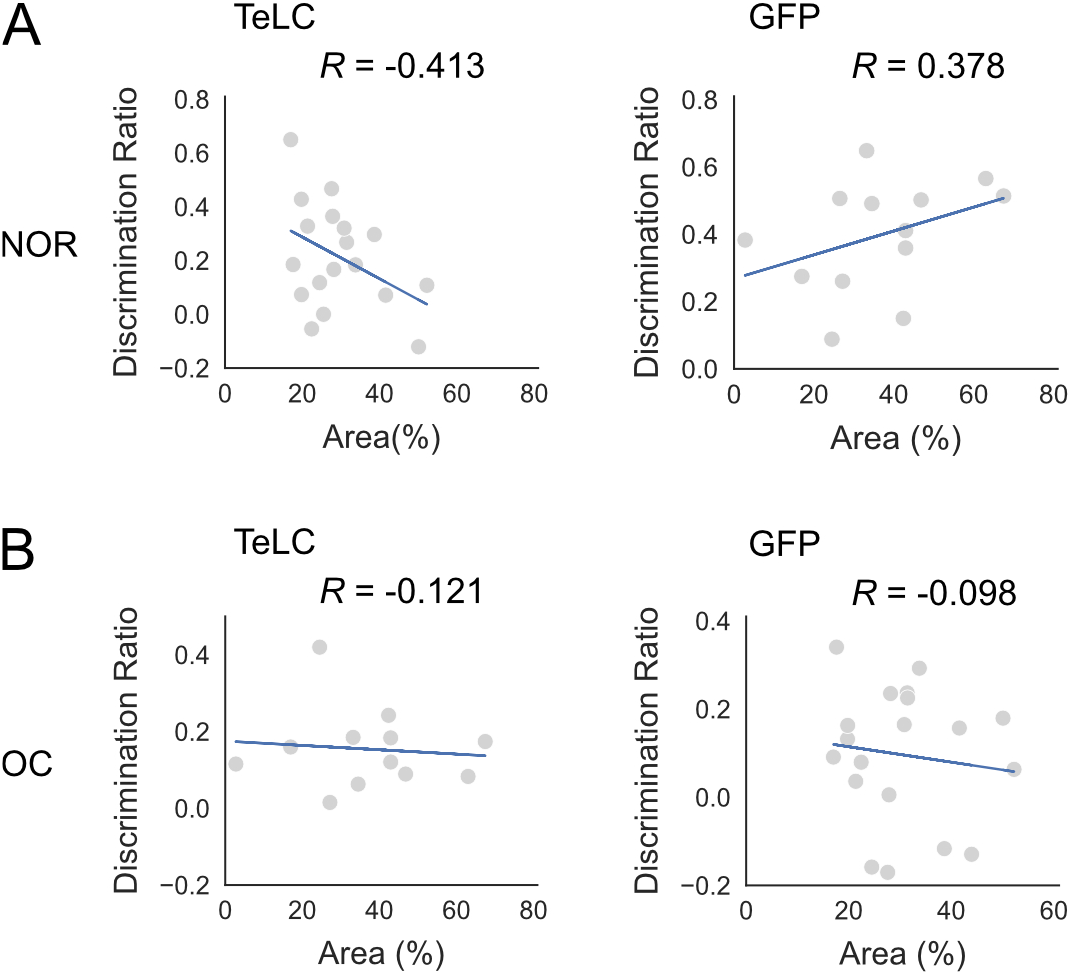
Relationship between virus expression and performance for novel object recognition (NOR) and object-context (OC) tasks. Scatterplots of the percentage area of virus expression in L2 plotted against the discrimination ratios for the NOR (A) and OC task (B). Each grey dot represents data for a single animal. Pearson’s correlation coefficients (*R*) were calculated to determine the strength of the relationship between discrimination ratios and virus expression. The exact value of *R* is indicated in the top right corner of each plot. There was no significant correlation between discrimination ratios and virus expression in either group for the NOR (TeLC: *P* = 0.089; GFP control: *P* = 0.202) or OC task (TeLC: *P* = 0.622; GFP control: *P* = 0.762). Each plot is overlaid with line of best fit (blue) that was calculated using the least squares method of linear regression.

